# Multi-dimensional tuning in motor cortical neurons during active behavior

**DOI:** 10.1101/2020.04.03.022178

**Authors:** Rachel C. Yuan, Sarah W. Bottjer

## Abstract

A region within songbird cortex, AId (dorsal intermediate arcopallium), is functionally analogous to motor cortex in mammals and has been implicated in vocal learning during development. AId thus serves as a powerful model for investigating motor cortical contributions to developmental skill learning. We made extracellular recordings in AId of freely behaving juvenile zebra finches and evaluated neural activity during diverse motor behaviors throughout entire recording sessions, including song production as well as hopping, pecking, preening, fluff-ups, beak interactions with cage objects, scratching, and stretching. A large population of single neurons showed significant modulation of activity during singing relative to quiescence. In addition, AId neurons demonstrated heterogeneous response patterns that were evoked during multiple movements, with single neurons often demonstrating excitation during one movement type and suppression during another. Lesions of AId do not disrupt vocal motor output or impair generic movements, suggesting that the responses observed during active behavior do not reflect direct motor drive. Consistent with this idea, we found that some AId neurons showed differential activity during pecking movements depending on the context in which pecks occurred, suggesting that AId circuitry encodes diverse inputs beyond generic motor parameters. Moreover, we found evidence of neurons that did not respond during discrete movements but were nonetheless modulated during active behavioral states compared to quiescence. Taken together, our results support the idea that AId neurons are involved in sensorimotor integration of external sensory inputs and/or internal feedback cues to help modulate goal-directed behaviors.

**SIGNIFICANCE STATEMENT:** Motor cortex across taxa receives highly integrated, multi-modal information and has been implicated in both execution and acquisition of complex motor skills, yet studies of motor cortex typically employ restricted behavioral paradigms that target select movement parameters, preventing wider assessment of the diverse sensorimotor factors that can affect motor cortical activity. Recording in AId of freely behaving juvenile songbirds that are actively engaged in sensorimotor learning offers unique advantages for elucidating the functional role of motor cortical neurons. The results demonstrate that a diverse array of factors modulate motor cortical activity and lay important groundwork for future investigations of how multi-modal information is integrated in motor cortical regions to contribute to learning and execution of complex motor skills.

## INTRODUCTION

Goal-directed motor learning underlies our ability to both acquire new motor skills and flexibly perform those skills in response to changing environmental contexts. Increasing evidence indicates that motor cortex serves as not only a driver of learned movements but also a central locus for motor learning. For instance, skill learning, disease or trauma can induce experience-dependent changes in motor cortical representations, including expansions of motor maps that may reflect functional reorganization to support newly learned movements (Sanes and Donoghue, 2000; Li et al., 2001; Conner et al., 2005; Xu et al., 2009; Makino et al., 2016; Peters et al., 2017; Papale and Hooks, 2018). Moreover, while the necessity of motor cortex for performance of learned skills can vary depending on factors such as the level of dexterity required or the amount of time spent training, motor cortex is required for the acquisition of new motor skills across a variety of movements and training paradigms (Whishaw et al., 1991; Whishaw, 2000; Darling et al., 2011; Guo et al., 2015; Kawai et al., 2015; Hwang et al., 2019).

Both learning and performance of motor skills are highly sensorimotor processes: successful acquisition of a motor behavior entails integration across internal and external sources of sensory feedback in order to guide accurate refinement of motor output. Feedback information is also required to flexibly perform an acquired skill in response to changing environmental demands post-learning; even relatively simple movements such as grasping an object require integration across modalities in order to select the appropriate motor action for a given behavioral context. Correspondingly, motor cortical neurons demonstrate multi-dimensional tuning that suggests integration across a variety of sensorimotor aspects: motor cortical neurons involved in performance of a skill have been shown to encode not only motor parameters (e.g., movement speed or direction) but also non-motor parameters, such as preparatory activity prior to movement execution or visual information specific to a target object’s location in object-directed reaching tasks (Tanji and Evarts, 1976; Evarts and Fromm, 1977; Murata et al., 1997; Shen and Alexander, 1997). Taken together, these findings suggest motor cortex as a dynamic substrate that actively integrates diverse streams of information to contribute to sensorimotor learning and performance. However, the potential influence of various multi-modal inputs on motor cortical activity during behavior is difficult to assess in anesthetized and/or restrained experimental paradigms that focus on a single motor task; recordings in freely behaving animals, especially in the context of sensorimotor skill learning, provide an attractive alternative for investigating sensorimotor integration in motor cortical neurons (Ebbesen et al., 2017; Zhang et al., 2017; Mimica et al., 2018).

Vocal learning in songbirds entails integration of social cues and visual, auditory, and somatosensory feedback information to guide refinement of variable juvenile babbling into stereotyped song and thus serves as a powerful model for investigating goal-directed sensorimotor learning (Price, 1979; West and King, 1988; Eales, 1989; Mann et al., 1991; Mann and Slater, 1995; King et al., 2005; Derégnaucourt et al., 2013; Chen et al., 2016; Ljubičić et al., 2016; Carouso-Peck and Goldstein, 2019). In songbirds, the cortical region AId lies within an area that has been considered to be analogous to motor cortex in mammals: akin to infragranular layers of mammalian motor cortex, AId receives inputs that process multi-modal sensory information via dNCL (dorsal caudolateral nidopallium) as well as information from cortico-basal ganglia circuitry that is dedicated to vocal learning, and in turn makes a variety of projections that give rise to feedforward and feedback pathways through subcortical and brainstem regions (Fig. 1A) (Zeier and Karten, 1971; Bottjer et al., 2000; Karten, 2013; Paterson and Bottjer, 2017). Lesions of AId in juvenile zebra finches impair the bird’s ability to achieve an accurate imitation of its memorized tutor song but do not induce vocal motor deficits (Bottjer and Altenau, 2010), suggesting an important role for this region in guiding motor refinement during sensorimotor learning. AId is thus well suited for investigation of motor cortical contributions to goal-directed acquisition of complex motor skills.

**Figure 1.**
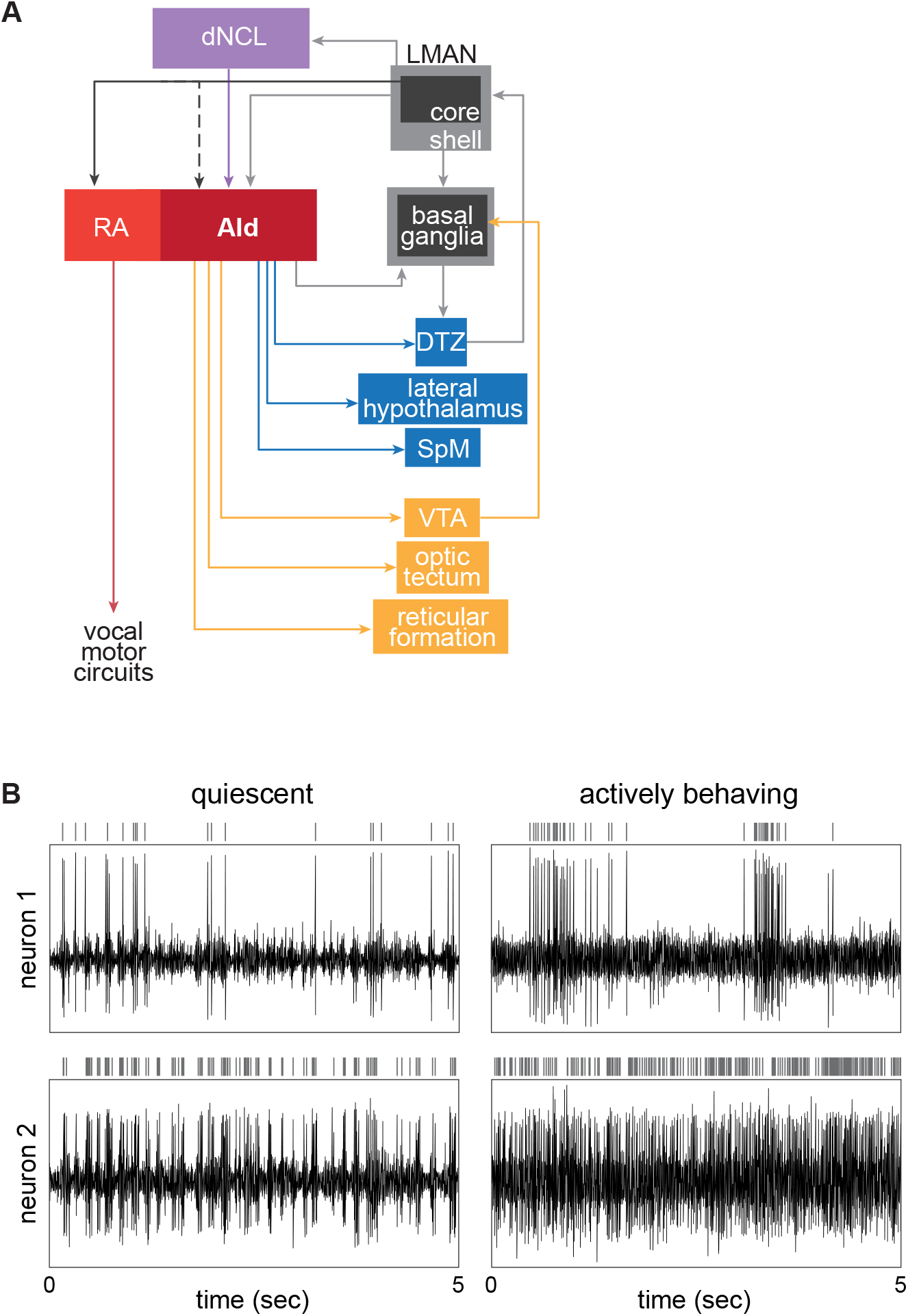
AId neurons are well situated to integrate multi-modal inputs and distribute information across various cortical-subcortical circuits. ***A***, AId is a motor cortical region that lies within an arcopallial area that has been considered to be analogous to motor cortex in mammals. AId receives inputs from upstream cortical regions LMAN-shell and dNCL. LMAN-shell is part of a cortico-basal ganglia loop that mediates vocal learning, whereas dNCL receives inputs from LMAN-shell as well as multiple pathways processing somatosensory, visual, and auditory information. AId of juvenile birds also receives inputs from LMAN-CORE via axon collaterals of LMAN-CORE→RA neurons that drive vocal output; robust collaterals are present in juvenile birds prior to ∼40-45 dph but are not present in older juvenile or adult birds. AId projects to striatum and several midbrain and thalamic regions that give rise to both feedback and feed-forward pathways, creating several opportunities for information transfer between cortical and subcortical regions. See text. Abbreviations: AId, dorsal intermediate arcopallium; dNCL, dorsal caudolateral nidopallium; DTZ, dorsal thalamic zone; LMAN, lateral magnocellular nucleus of the anterior nidopallium; RA, robust nucleus of the arcopallium; SpM, medial spiriform nucleus; VTA, ventral tegmental area. ***B***, Raw traces of extracellular activity simultaneously recorded at two different sites within AId of a juvenile bird (44 dph) while the bird was resting (left column; “quiescent”) versus hopping around the recording cage (right column; “actively behaving”). Vertical lines above each raw activity trace indicate spikes from a single neuron sorted from the extracellular activity.

To investigate the role of AId neurons in motor skill learning, we made extracellular recordings in AId of freely behaving juvenile songbirds during the sensorimotor period of vocal learning. In addition to recording AId neurons during singing behavior, we analyzed activity during several discrete movements and across different state periods based on the bird’s behavior. Our results thus represent an extensive assessment of motor cortical activity across a wide variety of natural behaviors, thereby informing our understanding of how these neurons may contribute to motor skill learning and production.

## MATERIALS AND METHODS

### Subjects

All animal procedures were performed in accordance with the University of Southern California animal care committee’s regulations. Seven male juvenile zebra finches (43-58 days post hatch, dph; mean age 46 dph on first day of recording) were used. All birds were raised in group aviaries until at least 33 dph, remaining with their natural parents and thereby receiving normal auditory and social experience during the tutor memorization period (Immelmann, 1969; Böhner, 1983, 1990; Eales, 1985; Mann and Slater, 1995; Roper and Zann, 2006). Juveniles were separated from group aviaries at 33-35 dph and housed in single cages within the experimental rig. Each bird’s tutor was placed in a separate cage within view of the juvenile to help it acclimate to the experimental rig for 2-5 days prior to the start of recording.

### Electrophysiology

At 35-40 dph, birds were anesthetized with 1.5% isoflurane (inhalation) and placed in a stereotaxic instrument. An electrode assembly consisting of four tetrodes affixed to a movable microdrive was fixed to the skull using C&B Metabond (Parkell), such that the tetrodes were implanted 500 μm dorsal to AId. Each tetrode consisted of four twisted polyimide-coated Nichrome wires (0.012 mm diameter Redi Ohm 800, Kanthal) routed through fused silica capillary tubing and electroplated with non-cyanide gold plating solution (SIFCO 5355). One day after surgery, the tetrode assembly was connected to a recording headstage (HS-16, Neuralynx) with a flexible cable connected to a commutator (PSR, Neuralynx); 15 channels of neural data were amplified, band passed between 300 and 5000 Hz (Lynx-8, Neuralynx), and digitized at 32 kHz using Spike2 software (Power 1401 data acquisition interface, CED). Audio and video were recorded coincident with neural activity – vocalizations were recorded with a lavalier microphone (Sanken COS-11D) mounted in the cage; a USB-video camera (30 FPS, ELP) was placed at the front of the cage to record video. Consecutive 30-minute recordings were made from 7:00 AM to 6:00 PM each day. Tetrodes were manually advanced with the microdrive when the cells being recorded were lost or had already been recorded for at least two days, as indicated by stability and consistency of the extracellular signal. At the end of each experiment, birds were perfused (0.7% saline followed by 10% formalin), and brains were removed and post-fixed for 72 hours before being cryo-protected (30% sucrose solution) and frozen-sectioned (50 μm thick). Sections were Nissl-stained with thionin to visualize tetrode tracks and verify recording locations.

Movement artifact in neural recordings was correlated across recording channels and was eliminated or reduced using offline common average referencing: for each recording channel, the signal across ∼8-14 remaining recording channels was averaged and subtracted from that channel in order to remove movement artifact (Ludwig et al., 2009). Channels were visually inspected after referencing to ensure that spiking activity was not distorted. After common average reference subtraction, single units were sorted from multi-unit data by first automatically clustering units with KlustaKwik (KD Harris, University College London). KlustaKwik clusters were manually inspected across 18 different waveform features and further refined using MClust (AD Redish, University of Minnesota). Clusters were considered for analysis only if the signal to noise ratio was > 2 and less than 1% of spikes had an inter-spike interval < 2 ms.

### Behavioral scoring

Sixteen 30-minute sessions were recorded across seven birds (median of 3 sessions per bird). Videos from recording session were scored for movements and state periods using ELAN (The Language Archive, Max Planck Institute for Psycholinguistics). We scored each single occurrence of pecks, hops, preening behavior, beak interactions (beak wipes and periods when the bird’s beak was in contact with perches, food cup edges, etc for longer than the duration of a peck), fluff-ups, scratches, and stretches. Head movements occurred so frequently that it was impractical to score all of them for all cells. However, we scored head movements that occurred during singing and during 30 seconds of non-singing before and after each singing episode in a subset of 36 singing-responsive neurons. These head movements were scored in order to compare the responses of neurons during head movements that occurred within singing versus non-singing periods. For 12 neurons across two recording sessions, all head and postural movements across the entire session were also scored.

In addition to scoring discrete movements, we developed a novel way of measuring behavior throughout recording sessions: each session was segmented into contiguous time periods that were classified into one of five behavioral “state” periods based on the bird’s behavior: eating, singing, active-movement, quiet-attentive, or quiescent; these state periods tiled the entire duration of the recording session (Fig. 6A). Eating states were defined as periods during which the bird was pecking at seed or grit, hulling or ingesting seeds, or pausing in between these behaviors for at most one second. Although pecks occurred most often during eating, eating states could also include other scored movements such as hops or preening, or unscored movements such as head movements as long as they occurred during the brief (≤ 1 sec) pauses that occurred while birds were actively eating. Singing states were defined by song behavior; they began whenever the bird produced song and lasted as long as song syllables continued to occur within one second of each other (inter-syllable interval ≤ 1 sec). Birds often made head movements during singing and occasionally made scored movements such as hopping or pecking in between bouts of singing. Active-movement states were defined as non-eating and non-singing periods during which the bird made active movements with pauses of at most one second in between movements; these periods could include any of the seven movements that were scored as well as head and postural body movements that were not scored. Quiet-attentive states were defined by times when the bird was not eating, singing, or moving around the cage for more than one second; they continued as long as the bird made at most small head movements and otherwise remained unmoving but alert. Quiescent periods were defined as periods during which the bird was completely still and not obviously paying attention to any stimulus, with eyes partially or fully closed. Quiescent state times were segmented into one-second intervals that were used as baseline periods for analyses of scored movements (see below).

### Data Analysis

To test for significant responses during scored movements, firing rates across occurrences of each movement type were compared against quiescence. Quiescent baseline periods were generated by dividing quiescent state periods (as described above) into one-second segments. The firing rate during two one-second quiescent segments that occurred closest in time to each movement occurrence was used as a corresponding baseline. Fourteen neurons were recorded during sessions that lacked quiescent state periods. For these 14 neurons, one-second baseline periods were taken from times within quiet-attentive state periods when the bird was verified to be unmoving (though clearly alert, unlike in quiescent state). To compare movement responses across neurons, standardized response strength was calculated for each movement type as:

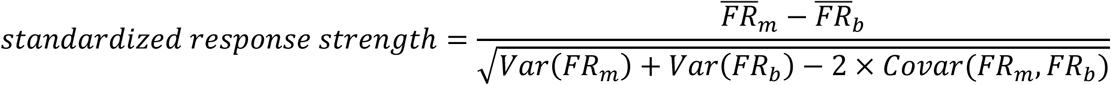

where FR_m_ is the firing rate during movement occurrences and FR_b_ is the firing rate during corresponding baseline periods. A positive value indicates an increase in firing rate during the movement compared to quiescence whereas a negative value indicates a decrease in firing rate during movement. This measure is referred to as response strength (RS) throughout the text.

To test for changes in activity around movement onsets, for each neuron we generated a 25 ms-bin histogram of the spiking response across all occurrences of the movement; histogram windows were one second long and centered on movement onsets. Spike times during each movement repetition were shuffled to obtain a resulting histogram of shuffled spike data; this was repeated 1000 times, resulting in 1000 histograms of shuffled data. Each bin of the actual spike data histogram was considered significantly excited if the count in that bin was greater than 95% of maximum values from the shuffled data set; likewise, the bin was considered significantly suppressed if the count was lower than 95% of minimum values from the shuffled data set. Onset responses were defined as responses that contained at least two consecutive bins (50 ms) of significant maxima or minima within 100 ms of movement onset.

We tested for significant offset-aligned responses using the same method and parameters as onset-aligned responses, except that the one-second windows used for histograms of actual and shuffled data were centered around movement offsets; offset responses were defined as responses that contained at least two consecutive bins (50 ms) of significant maxima or minima within 100 ms of movement offset. Due to the short duration of pecks and hops (mean hop duration = .28 seconds; mean peck duration = .25 seconds), it was possible for the same maxima or minima to be captured in both onset- and offset-aligned responses. To ensure that offset-aligned activity could be accurately distinguished, for these movements we only counted excited or suppressed offset responses that did not demonstrate significant changes in onset-aligned activity. Preening movements were relatively long in duration (mean duration = 1.5 seconds), so all preening-aligned offset responses were counted.

We tested for significant modulation of firing rate during states by dividing all state periods in each recording session into one-second segments and calculating the average firing rate during each segment for each neuron. We compared the distributions of average firing rates across segments for each state against quiescence to determine whether activity was significantly increased or decreased during non-quiescent states for each neuron. Fourteen neurons were recorded during sessions that lacked quiescent state periods and were excluded from these analyses.

As an additional means of characterizing firing rate modulation during states, we defined “events” – brief periods of excitation and suppression – from histograms of spiking activity. We segmented the instantaneous firing rate (IFR) across each recording session into 10-ms bins and smoothed the IFR with a moving average filter (span = 3 bins). Excited events were defined as periods during which the smoothed IFR across five or more 10-ms bins (50 ms or more) exceeded the average firing rate across quiescent state periods by ≥ 3 standard deviations. Suppressed events were defined as periods during which the smoothed IFR across five or more 50-ms bins fell below the average firing rate across quiescent state periods by ≤ 1.5 standard deviations. To compare across neurons, the number of events in each state was normalized by dividing the number of excited or suppressed events in each state by the total duration of that state for each neuron.

### Statistics

Movement responses were tested for significance against quiescent baselines (see above) using Wilcoxon signed rank tests; Benjamini-Hochberg post-hoc tests were used to apply corrections for multiple comparisons (Benjamini and Hochberg, 1995). Neurons that demonstrated a significant difference between movement and baseline for at least one scored movement were considered movement-responsive. To test whether movement responses were context-selective, response strengths during movements from one context versus another context were compared using Mann-Whitney tests for each neuron (for example, comparing pecks during eating versus non-eating periods, and comparing head movements during singing versus non-singing periods). Mann-Whitney tests were also used to compare response strengths during singing periods with versus without head movements in individual neurons. χ^2^ tests were run to compare proportions between more than two groups (for example, proportions of neurons that were responsive during each movement type). In case of significance, Fisher’s exact tests were used as a follow-up to make pairwise comparisons of proportions between groups, and Benjamini-Hochberg post-hoc tests were used to apply a correction for multiple comparisons. Binomial tests were used to judge whether the relative proportions of excited versus suppressed responses among movement-responsive neurons were different from chance. Comparisons of firing rate distributions between each state and quiescence (see above) were made using Kolmogorov-Smirnov tests, with a Benjamini-Hochberg post-hoc test applied for multiple comparisons. Measures between different state periods (inter-spike intervals, normalized number of events, normalized firing rates among unresponsive neurons) were compared using sets of pairwise linear contrasts based on trimmed means (20% trimming); this linear contrast method has been shown to be robust to common assumption violations such as non-normality and heteroscedasticity (Wilcox and Serang, 2017). For all tests, p < .05 was considered significant.

## RESULTS

We made extracellular recordings from 119 neurons in AId of freely behaving juvenile zebra finches (43-58 dph) housed singly in a recording cage as they actively engaged in sensorimotor vocal practice. A typical 30-minute recording period included various overt behaviors and periods of quiescence when the bird was not moving. To investigate how neural activity in AId corresponds to different behaviors, we scored seven different movements during each recording that could be identified reliably: pecks, hops, preening episodes, beak interactions with objects in the recording cage (e.g., beak wipes or non-peck interactions with cage bars), fluff-ups, stretches, and scratching episodes; we also marked periods of singing. We developed a novel approach in which we examined spiking patterns of single neurons throughout each recording period to investigate whether the activity of AId neurons is selective for different movement types and/or singing behavior in juvenile birds.

### Responsivity of AId neurons during movements

Patterns of spiking were highly variable across individual neurons, ranging from phasic to tonic activity. In addition, each neuron’s activity was highly modulated throughout a typical recording session, showing either excitation and/or suppression during different movements. Figure 1B illustrates two different neurons recorded in one bird while it was quiescent (no overt movements, left columns) and while it was hopping around the cage (right columns). The neuron in the top panel fires intermittently in small bursts of at most three spikes during quiescence, while the neuron in the bottom panel exhibits dense bursting activity. As the bird hops around the cage, the neuron in the top panel shifts to longer periods of high firing separated by relative inactivity while the neuron in the bottom panel shifts to a high tonic rate of firing. To investigate whether such modulations were related to specific movements, for each neuron we compared firing rates during different movement types against baseline firing rates during quiescent periods that were closest in time to each movement occurrence (see Materials and Methods). To compare movement-related activity across neurons, we also calculated the response strength of each neuron during each movement type, defined as the standardized difference in average firing rate during each movement type versus baseline periods (see Materials and Methods).

The top plot in Figure 2A shows that 101 out of 119 neurons (85%) exhibited a significant change in firing rate during at least one movement type compared to quiescent baseline periods and were thus classified as “movement responsive.” Among movement-responsive neurons, 33 out of 101 (33%) responded during only one scored movement whereas 68 out of 101 (67%) responded during two or more movements (Fig. 2A, bottom). Few cells responded during five or six movements, and no cells responded during all seven movements. Figure 2B depicts patterns of responsivity in these 101 neurons. For instance, 10 neurons were either excited or suppressed during pecks but were otherwise not responsive during any of the other six scored movements; likewise, two other groups of 10 neurons each selectively modulated their firing rate only during preening or hops. Neurons that responded during two movements could show either excitation or suppression during both movements, or excitation during one and suppression during the other. In general, responsivity across cells was heterogeneous – for instance, the subpopulation of neurons responsive during four movements showed excitation or suppression during various combinations of movements that could include any four of the seven scored behaviors.

**Figure 2.**
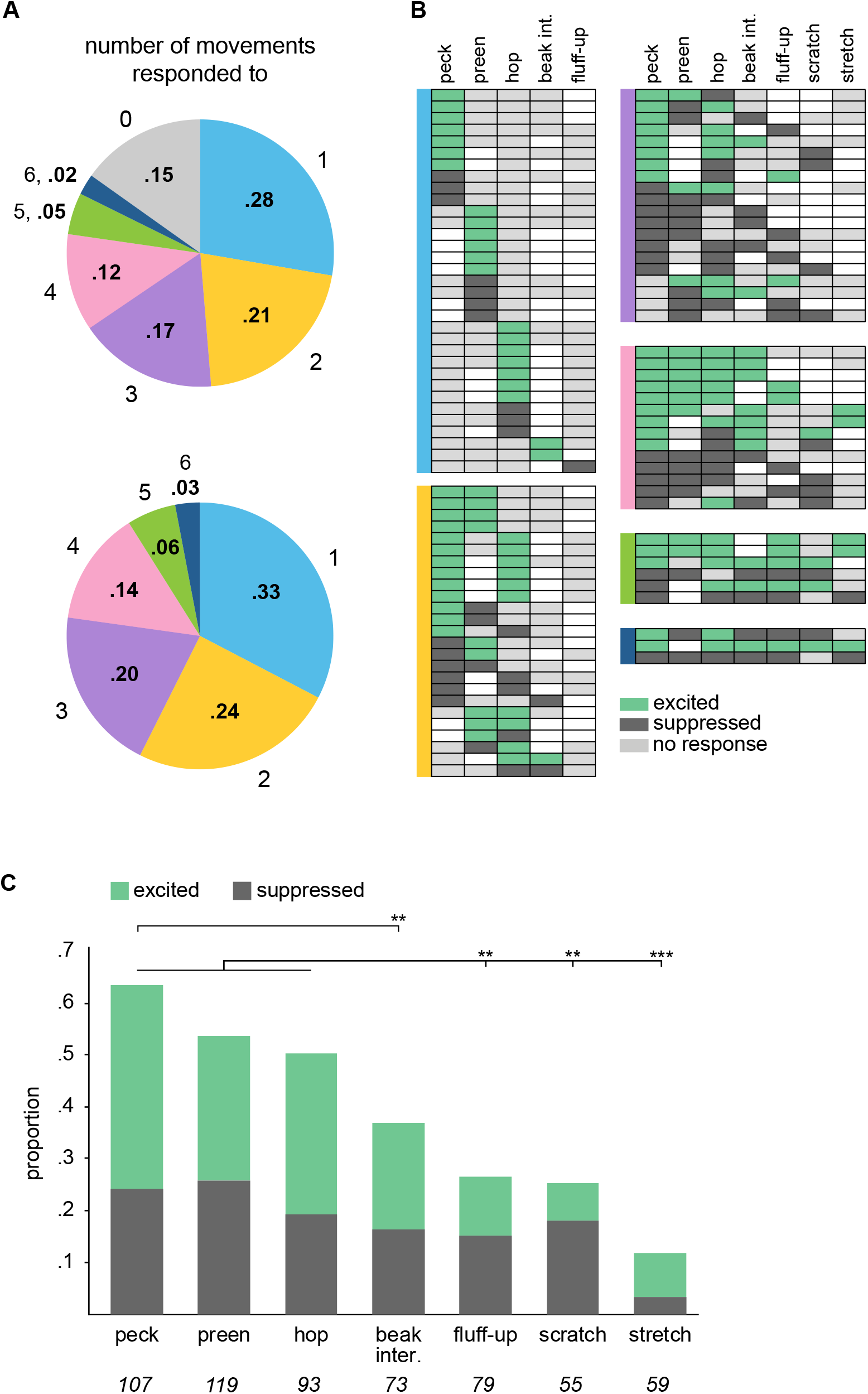
AId neurons respond during different scored movements with excitation and/or suppression. ***A***, Proportions of single AId neurons that responded during different numbers of movements. Top: proportions out of 119 neurons recorded in AId; 18/119 neurons were not movement responsive, while 33/119 neurons responded during one movement, 25/119 during two, 20/119 during three, 14/119 during four, 6/119 during five, and 3/119 during six movements. Bottom: proportions out of 101 AId neurons that responded during at least one movement. ***B***, Each row of each chart indicates movements during which each neuron was excited (green), suppressed (dark gray), or not responsive (light gray). Unfilled boxes indicate that no data during that movement was recorded for that neuron. Charts are grouped according to colors in ***A***, based on the number of movements during which neurons responded. ***C***, Proportions of AId neurons that were significantly excited (green) or suppressed (dark gray) during each movement type. Italicized numbers indicate the number of neurons recorded during each movement type. \*\**p*<.005, \*\*\**p*<.0001.

Higher proportions of neurons showed altered firing rates during pecks, preening, and/or hops compared to other movements: 68 out of 107 neurons (64%) were significantly modulated during pecks, 50 out of 93 (54%) during preening, and 60 out of 119 (50%) during hops (Fig. 2C). These proportions did not differ (Fisher’s exact test, Benjamini-Hochberg corrected, peck versus preen p = .23, peck versus hop p = .09, preen versus hop p = .71), and were each greater than the proportions of neurons that responded during fluff-ups (21/79, 27%), scratches (14/55, 25%), and stretches (7/59, 12%) (Fig. 2C; Fisher’s exact test, Benjamini-Hochberg corrected, p < .05 for all comparisons between pecks, hops, and preening to each of the other three movements; p > .05 for all comparisons among these latter three movements). In addition to the seven scored movements, birds constantly made quick, saccade-like movements throughout recording periods, easily resulting in over a thousand head or postural movements in a typical 30-minute session. As an initial test of whether firing rate was modulated relative to these movements, we scored all head and postural movements in a subset of 12 neurons and found that six neurons were excited during these movements (50%) while four were suppressed (33%), demonstrating that AId activity can be modulated during head and postural movements as well.

As indicated above, we observed both excited and suppressed responses within single neurons: 21 out of 68 neurons (31%) that responded during multiple movements exhibited excitation during some movements and suppression during others (Fig. 2B). The proportion of excited versus suppressed responses across all movement responses did not differ (56%, 138/247 excited; 44%, 109/247 suppressed; binomial test, p = .07), indicating a fairly even representation of excitation and suppression across scored movements. In addition, within each group of neurons that were responsive during a given movement, the proportions of excited versus suppressed responses did not differ (Fig. 2C; binomial test, p > .05 in all cases).

Overall, these results demonstrate that the responsivity of AId neurons is highly heterogeneous. Neurons modulated their firing rate most often during pecks, hops, and/or preening but varied in their responsivity profile: approximately half of the recorded neurons responded during only one or two movements, whereas the remainder were more broadly responsive and showed significant excitation or suppression during different combinations of movements.

### Temporal response patterns of AId neurons at movement onsets and offsets

Some AId neurons demonstrated consistent temporal changes in firing rate at movement onsets and/or offsets that could be masked by measures of average firing rate. For example, Figure 3A shows rasters and histograms for a single neuron during preening (left) and peck (right) responses. In both cases, mean firing rate during the movement was significantly excited relative to quiescence (preening RS = 0.65, peck RS = 1.53, p < 0.05 in both cases). However, the responses during both movements clearly contain periods of suppression that begin prior to movement onset as well.

**Figure 3.**
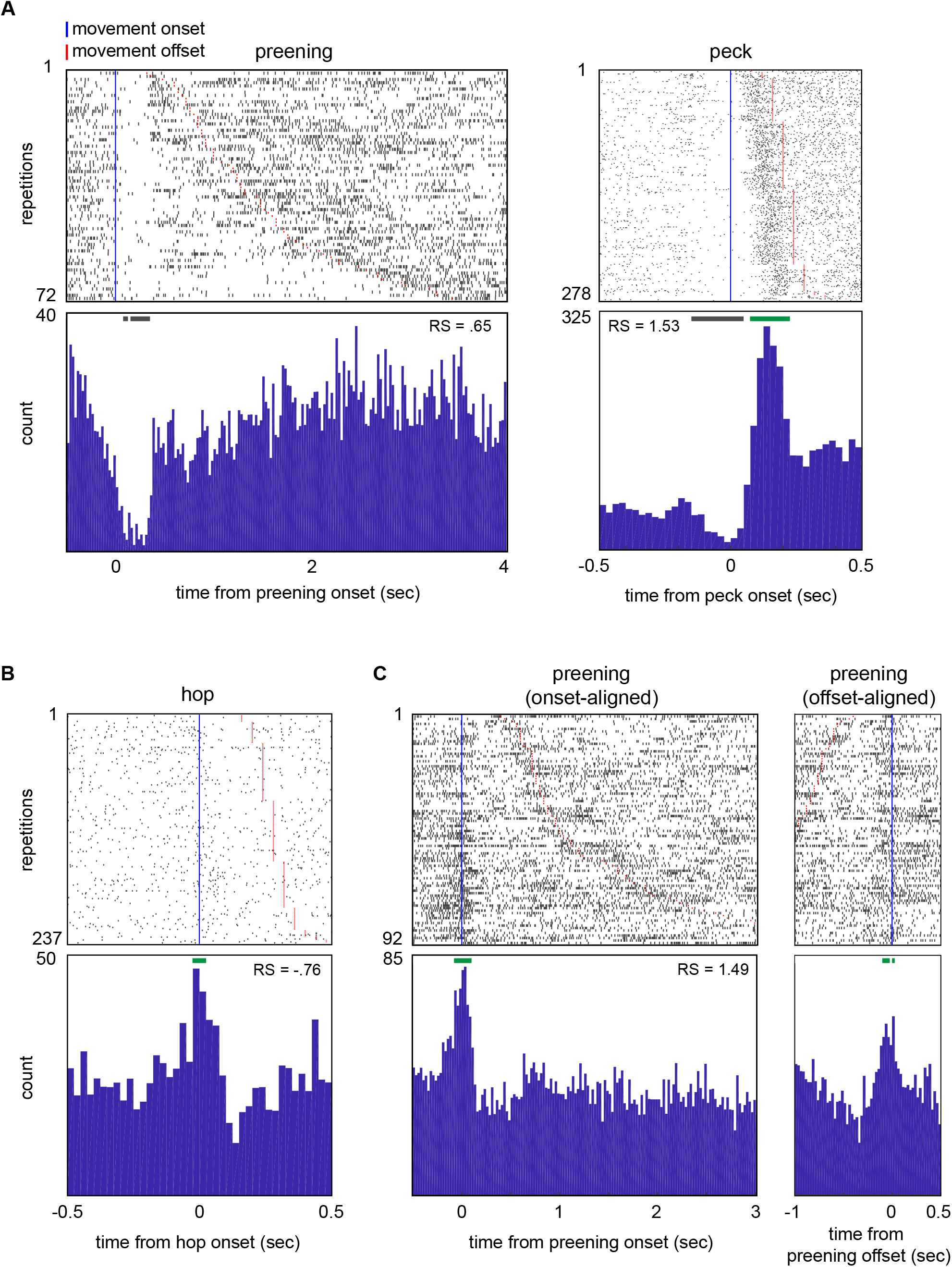
AId neurons show a variety of temporal response patterns during different scored movements. ***A***, Rasters and histograms illustrating the response of a single AId neuron during preening (left) and pecks (right). ***B***, Raster and histogram illustrating the response of a single AId neuron during hops. ***C***, Rasters and histograms illustrating the onset- (left) and offset-aligned (right) preening response of an example AId neuron. Rows are sorted by movement duration. Blue vertical lines mark movement onset; red lines mark movement offsets. Green and gray horizontal bars above histograms denote periods of excitation or suppression, respectively (see Materials and Methods). RS, average standardized response strength over the entire duration of each movement type.

To capture these firing rate modulations, we tested for periods of significant excitation or suppression at movement onsets and offsets. For each response, we compared histograms of spiking activity spanning one second centered on movement onsets or offsets to histograms of shuffled spike trains (25-ms bins) in order to identify bins in which the firing rate was significantly excited or suppressed (see Materials and Methods). Figure 3 plots various examples of onset-aligned (Fig. 3A, B; C, left) and offset-aligned (Fig. 3C, right) responses; green and gray horizontal bars above the histograms mark excited and suppressed bins, respectively. Onset or offset responses were defined as responses that included at least two contiguous bins (50 ms) of significant excitation or suppression occurring within 100 ms of movement onsets or offsets.

We observed significant onset responses during pecking, hopping, and preening, but not other movement types. The top of Table 1 lists onset responses for these three movements, classified by whether neurons showed a significant response based on average firing rate. Eleven out of 107 neurons (10.3%) exhibited significant excitation (7/11) or suppression (4/11) at peck onsets. Five out of 119 neurons demonstrated an onset response during hopping (4.2%; 4/5 excitation, 1/5 suppression), as did four out of 93 neurons during preening (4.3%; 3/4 excitation, 1/4 suppression). We also observed significant offset responses during pecking and preening: six out of 107 neurons exhibited pecking offset responses (5.6%; 4/6 excitation, 2/6 suppression), and three out of 93 neurons showed preening offset responses (3.2%; all excitation) (Table 1, bottom).

**Table 1.** Proportions of onset and offset responses across all neurons for different movement types

|  | **Peck-onset responses**<br>**($\pm 100$ ms from peck onset)** |  | **Hop-onset responses**<br>**($\pm 100$ ms from hop onset)** |  | **Preening-onset responses**<br>**($\pm 100$ ms from preening onset)** |  |
| --- | --- | --- | --- | --- | --- | --- |
| **Response based on average firing rate** | Excited (7) | Suppressed (4) | Excited (4) | Suppressed (1) | Excited (3) | Suppressed (1) |
| Excited | .05 (5/107) | .04 (4/107) | .01 (1/119) | 0 | .03 (3/93) | .01 (1/93) |
| Not significant | .02 (2/107) | 0 | .02 (2/119) | 0 | 0 | 0 |
| Suppressed | 0 | 0 | .01 (1/119) | .01 (1/119) | 0 | 0 |

|  | **Peck-offset responses**<br>**($\pm 100$ ms from peck offset)** |  | **Preening-offset responses**<br>**($\pm 100$ ms from preening offset)** |  |
| --- | --- | --- | --- | --- |
| **Response based on average firing rate** | Excited (4) | Suppressed (2) | Excited (3) | Suppressed (0) |
| Excited | .04 (4/107) | .01 (1/107) | .01 (1/93) | 0 |
| Not significant | 0 | 0 | .01 (1/93) | 0 |
| Suppressed | 0 | .01 (1/107) | .01 (1/93) | 0 |

Onset responses (top) and offset responses (bottom) are shown separately (total n = 29 responses), categorized based on whether average firing rate during the movement showed significant excitation (Excited), suppression (Suppressed), or no response (Not significant).

Out of 29 total onset and offset responses, 13 showed temporally patterned excitation or suppression that did not match the average firing rate response. For example, four responses (two pecking, two hopping) included consistent excitation at movement onset even though the average firing rate during movement was not significantly different from quiescent baseline (Table 1, top, “Not significant” row). Similarly, one preening response demonstrated significant offset-excitation but lacked a change in average firing rate across the entire movement (Table 1, bottom, “Not significant” row). Furthermore, onset and offset responses could differ in sign (excitation or suppression) from responses based on average firing rate. For example, while average firing rate during the hop response plotted in Figure 3B was suppressed relative to baseline (RS = -.76), the raster and histogram reveal an excitatory peak starting just before hop onset, indicating a complex temporal response that included brief excitation followed by suppression. Likewise, five neurons with excited average responses showed suppression at movement onset (four pecking, one preening; e.g., Fig. 3A; Table 1, top); one peck-excited response included suppression at movement offset, and one preening-suppressed response exhibited offset excitation (Table 1, bottom). These results suggest that single AId neurons can be modulated by multiple factors during movements, resulting in excitation at movement onsets or offsets and suppression during the movement itself, or vice versa.

### Context-dependency of pecking behavior

Of the seven movements we scored, pecking behavior in particular tended to occur in different contexts: birds always pecked while eating, but also pecked frequently at other objects such as cage bars or perches. To investigate whether such context influenced responsivity, we compared response strengths for pecks that occurred during eating versus other active behaviors (non-eating). Absolute values of response strength differed for pecks that occurred during eating versus non-eating in 47 out of 97 neurons (48%; Mann-Whitney test, p < .05). Figure 4A plots these context-sensitive cells according to whether they exhibited greater absolute response strength during eating (29/47, 62%; left) or non-eating (18/47, 38%; right).

**Figure 4.**
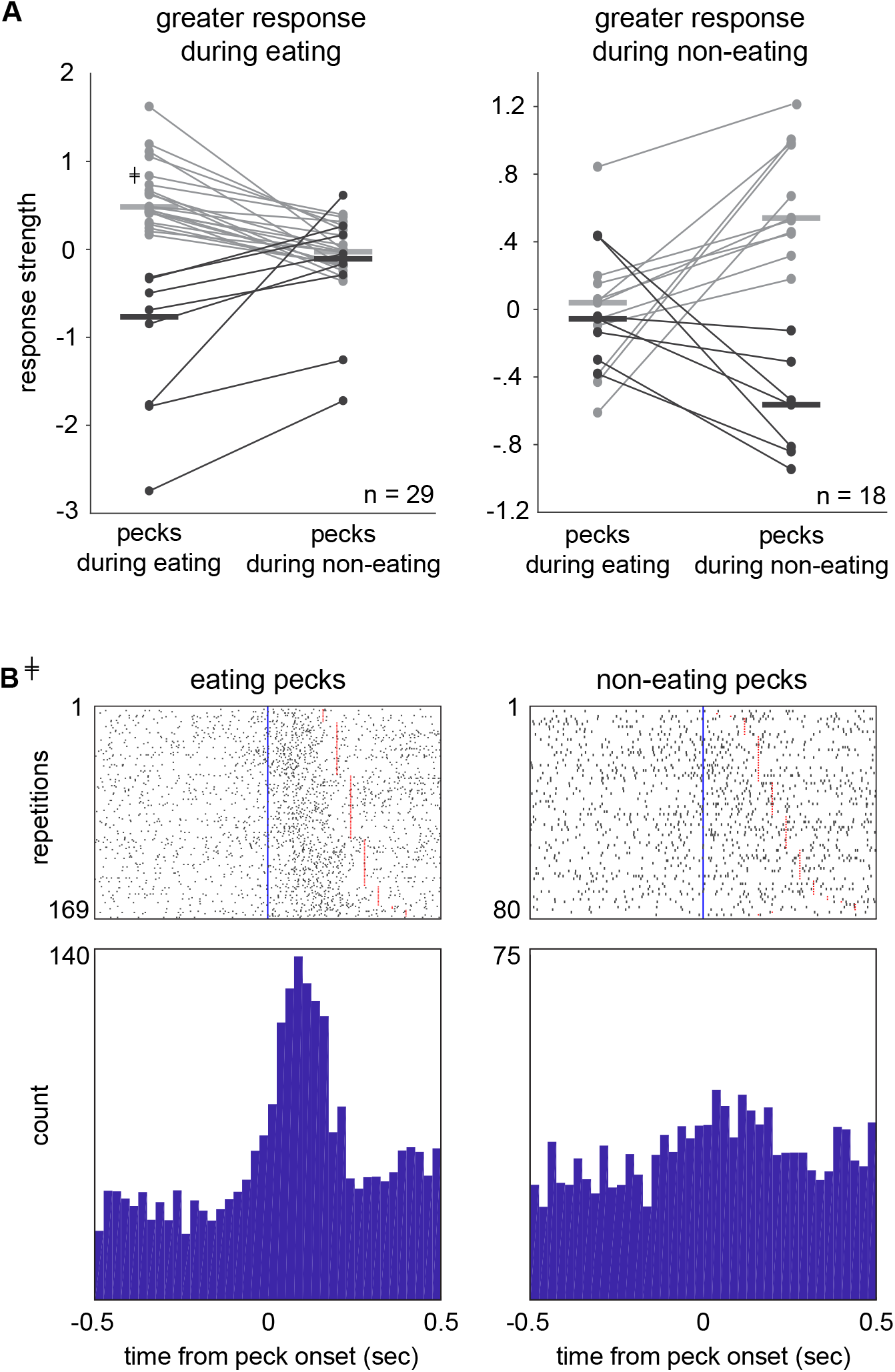
AId neurons exhibit context-sensitive peck responses. ***A***, Mean standardized response strengths of neurons during pecks that occurred during eating versus non-eating periods, grouped by neurons that showed greater absolute response strength during eating (left) and non-eating (right). Left: Gray and black lines represent neurons that showed positive or negative response strength, respectively, during eating-pecks. Right: Gray and black lines represent neurons that showed positive or negative response strength, respectively, during non-eating pecks. Response strengths during eating-related pecks were significantly different from non-eating pecks for all plotted neurons (Mann-Whitney test). ***B***, Rasters and histograms of an example neuron’s response during pecks that occurred during eating (left) versus non-eating (right). Peck response strength of this neuron is indicated by the cross-marked plot point in ***A***, left. Rows are sorted by peck duration. Blue vertical lines mark peck onsets; red lines mark peck offsets.

Among neurons that showed greater absolute response strength during eating-related pecks (mean absolute RS for eating-related pecks = .75 ± .11, non-eating pecks = .29 ± .07), most exhibited higher firing rates during eating-related pecks compared to non-eating pecks (21/29, 72%; Fig. 4A left, gray lines). For example, Figure 4B plots the peck-aligned responses for one of these neurons, illustrating strong excitation during pecks that occurred when the bird was eating (left) versus a lack of response during non-eating (right). However, eight out of 29 neurons (28%) showed lower firing rates during eating-related pecks compared to non-eating pecks, suggesting that AId neurons can signal different behavioral contexts with either relative excitation or suppression (Fig. 4A, left, black lines). In contrast to these “eating-peck” neurons, 18 neurons showed greater absolute response strength during pecks that were not associated with eating (mean absolute RS for eating-related pecks = .26 ± .05, non-eating pecks = .64 ± .07): 11 out of 18 neurons (61%) showed greater relative excitation for pecks during non-eating periods compared to eating periods (Fig. 4A, right, gray lines), whereas seven out of 18 (39%) showed greater relative suppression during non-eating pecks (Fig. 4A, right, black lines).

Overall, these results illustrate that pecking activity in many neurons was dependent on the context in which the movement occurred, suggesting that changes in response strength of these neurons is not specific to the physical movements of pecking behavior per se. One possibility is that these neurons are involved in processing orofacial or external sensory information that is present specifically in one context versus another.

### Singing-responsive neurons in AId

One of AId’s primary sources of afferents is from LMAN-shell, which contains neurons that are active during singing behavior and have been implicated in playing an important role in guiding accurate imitation of the tutor song during vocal learning (Fig. 1A) (Achiro and Bottjer, 2013; Achiro et al., 2017). Moreover, lesions of AId in juvenile birds impair the bird’s ability to achieve an accurate imitation of the adult tutor song without disrupting vocal motor output (Bottjer and Altenau, 2010). Given this evidence of an important role for AId in vocal learning, we hypothesized that firing rates of AId neurons would be modulated as juvenile birds were actively engaged in singing behavior.

As for analysis of individual movements, we compared firing rates of single neurons during song production against firing rates during quiescent baseline periods that were closest in proximity to each singing episode. Firing rates were significantly modulated during singing in 66 out of 94 neurons (70%), including 44 excited responses (mean RS = .76 ± .09) and 22 suppressed responses (mean RS = -.96 ± .19). Figure 5A illustrates the singing-aligned response of a neuron that increased its firing rate during song renditions. Altered firing rates during vocal production in songbirds have typically been interpreted as “singing-specific”. However, birds often make characteristic head and postural body movements during singing, as well as beak-gape and gular-fluttering movements that are specific to song production, raising the question of which movements are an intrinsic part of singing behavior versus independent movements that may be performed simultaneously during song production. Given the range of movement responsivity across AId neurons (Fig. 2), firing rate changes observed in singing-responsive neurons may reflect modulation by singing-specific actions as well as movements that are performed during both singing and non-singing periods. As an initial test of this question, we compared neural activity during discrete head movements that occurred within singing periods versus adjacent non-singing periods in a subset of 36 neurons that showed significant modulation of response strength during singing (see Materials and Methods).

**Figure 5.**
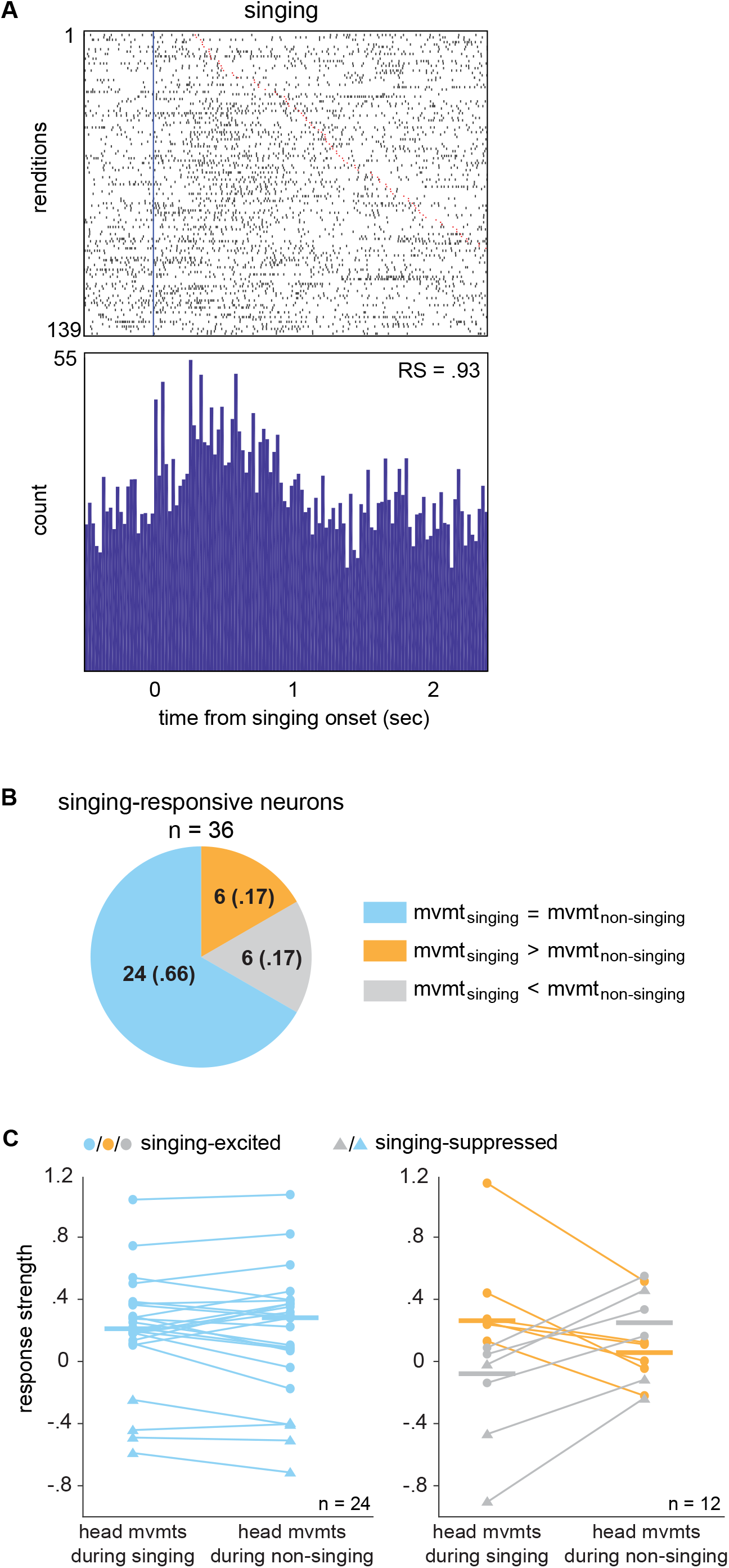
A substantial population of AId neurons are responsive during singing. ***A***, Raster and histogram illustrating activity of an example singing-excited AId neuron during singing episodes. Rows are sorted by duration of each singing episode. Blue vertical line marks onset of each singing episode; red lines mark ends of singing episodes. ***B***, Proportions of 36 singing-responsive neurons for which response strength during head movements that occurred within singing periods was greater than (orange), less than (gray), or not different from (blue) head movements that occurred during non-singing periods. ***C***, Mean standardized response strengths during head movements that occurred during singing versus non-singing periods. Lines connect data points from single neurons. Left: neurons that showed comparable response strength during head movements that occurred within singing versus non-singing periods. Right: neurons that showed higher response strength during head movements that occurred within singing (orange) or non-singing (gray) periods. Circles versus triangles represent neurons that showed an increase or decrease, respectively, in average firing rate across entire singing episodes relative to quiescence.

Response strength during head movements that occurred within singing periods did not differ from those within non-singing periods for most singing-responsive neurons (24/36, 67%; Mann-Whitney test, p > .05 for each neuron) (Fig. 5B; C, left). In eight of these 24 neurons, response strength during singing periods that included head movements was significantly greater than during singing periods that lacked head movements, indicating that activity during head movements contributed to the singing response (7/8 excited, 1/8 suppressed; Mann-Whitney test, p < .05 for each neuron; Table 2A, left). For each of the remaining 16 out of 24 neurons, response strength was modulated even during singing periods that lacked head movements and either did not differ from (14/24) or was greater than (2/24) the response during singing periods with head movements (Table 2A, middle and right). Thus, firing rate changes in most of these singing-responsive neurons was not attributable to activity during head movements.

**Table 2.**
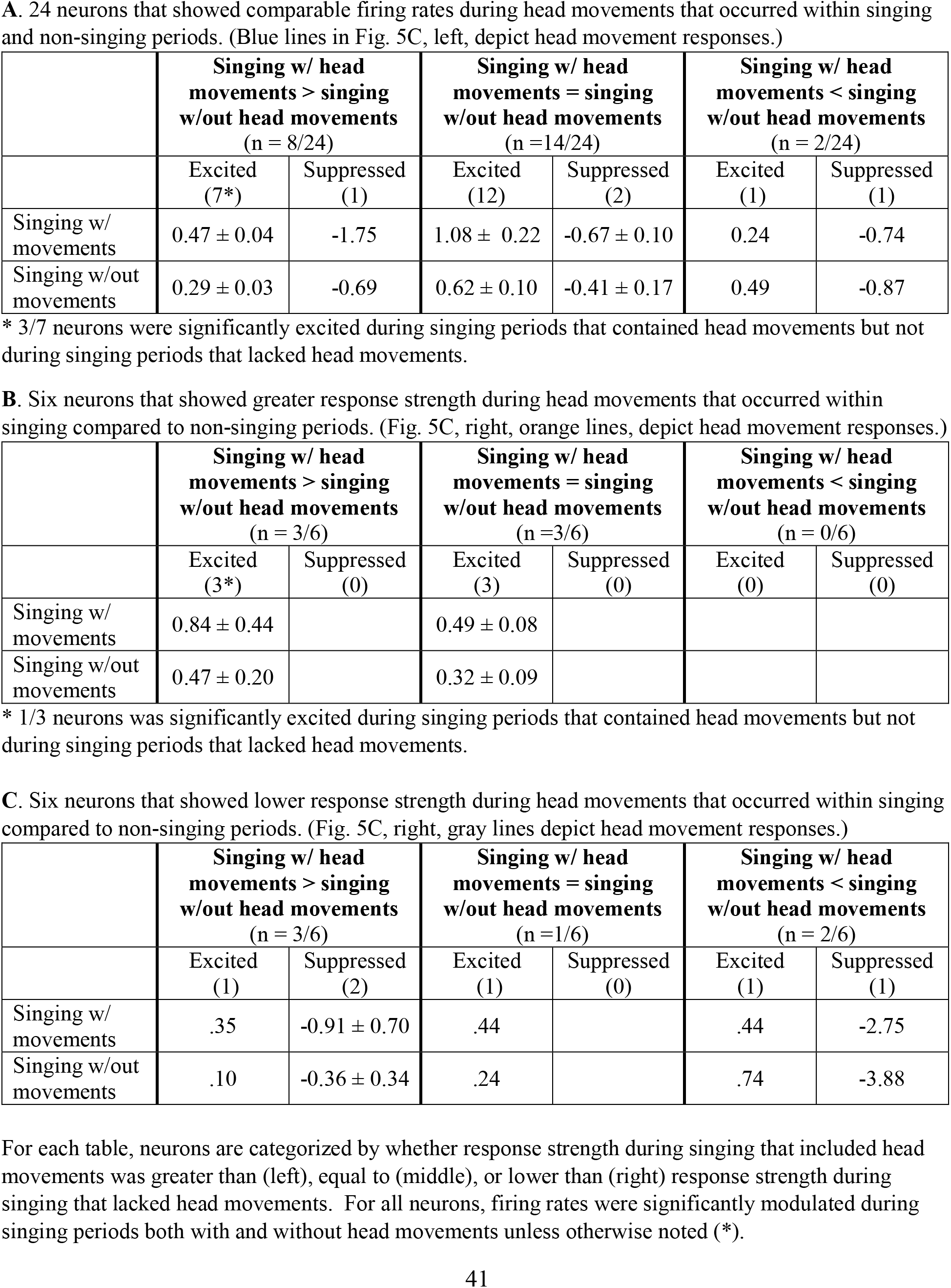
Mean standardized response strengths during singing periods with versus without head movements (n = 36 singing-responsive neurons)

|  | **Singing w/ head movements > singing w/out head movements**<br>(n = 8/24) |  | **Singing w/ head movements = singing w/out head movements**<br>(n = 14/24) |  | **Singing w/ head movements < singing w/out head movements**<br>(n = 2/24) |  |
| --- | --- | --- | --- | --- | --- | --- |
|  | Excited<br>(7*) | Suppressed<br>(1) | Excited<br>(12) | Suppressed<br>(2) | Excited<br>(1) | Suppressed<br>(1) |
| Singing w/ movements | 0.47 ± 0.04 | -1.75 | 1.08 ± 0.22 | -0.67 ± 0.10 | 0.24 | -0.74 |
| Singing w/out movements | 0.29 ± 0.03 | -0.69 | 0.62 ± 0.10 | -0.41 ± 0.17 | 0.49 | -0.87 |

\* 3/7 neurons were significantly excited during singing periods that contained head movements but not during singing periods that lacked head movements.

|  | **Singing w/ head movements > singing w/out head movements**<br>(n = 3/6) |  | **Singing w/ head movements = singing w/out head movements**<br>(n = 3/6) |  | **Singing w/ head movements < singing w/out head movements**<br>(n = 0/6) |  |
| --- | --- | --- | --- | --- | --- | --- |
|  | Excited<br>(3*) | Suppressed<br>(0) | Excited<br>(3) | Suppressed<br>(0) | Excited<br>(0) | Suppressed<br>(0) |
| Singing w/ movements | 0.84 ± 0.44 |  | 0.49 ± 0.08 |  |  |  |
| Singing w/out movements | 0.47 ± 0.20 |  | 0.32 ± 0.09 |  |  |  |

\* 1/3 neurons was significantly excited during singing periods that contained head movements but not during singing periods that lacked head movements.

|  | **Singing w/ head movements > singing w/out head movements**<br>(n = 3/6) |  | **Singing w/ head movements = singing w/out head movements**<br>(n = 1/6) |  | **Singing w/ head movements < singing w/out head movements**<br>(n = 2/6) |  |
| --- | --- | --- | --- | --- | --- | --- |
|  | Excited<br>(1) | Suppressed<br>(2) | Excited<br>(1) | Suppressed<br>(0) | Excited<br>(1) | Suppressed<br>(1) |
| Singing w/ movements | .35 | -0.91 ± 0.70 | .44 |  | .44 | -2.75 |
| Singing w/out movements | .10 | -0.36 ± 0.34 | .24 |  | .74 | -3.88 |

For each table, neurons are categorized by whether response strength during singing that included head movements was greater than (left), equal to (middle), or lower than (right) response strength during singing that lacked head movements. For all neurons, firing rates were significantly modulated during singing periods both with and without head movements unless otherwise noted (\*).

Six out of 36 neurons (17%) showed significant excitation during singing episodes and greater response strength during head movements that occurred within singing compared to non-singing periods (Fig. 5B; C, right, orange). In three of these neurons, activity during singing periods with head movements was comparable to singing periods without head movements, suggesting that discrete head movements made little contribution to changes in response strength during singing (Table 2B, middle). For the other three neurons, activity during singing periods with head movements was greater than during singing periods without head movements, indicating that singing-specific head movements contributed to excitation during song production (Table 2B, left). However, in all but one of these six neurons, activity during singing periods that lacked head movements was still significantly greater than quiescence.

Three neurons were singing-suppressed and showed lower response strength during singing-related head movements compared to non-singing (Fig. 5C, right, gray triangles). For two of these neurons, lower firing rates during head movements contributed to greater suppression during singing (Table 2C, left, Suppressed). However, all three neurons were suppressed even during singing periods that lacked head movements – in fact, for one of these neurons, suppression during singing periods without head movements was significantly greater than during periods with head movements (Table 2C, right, Suppressed). Interestingly, three neurons showed lower response strength during singing-related head movements compared to non-singing but nevertheless showed significant excitation across singing periods (Fig. 5C, right, gray circles), highlighting the presence of multiple modulating factors during song behavior.

In summary, most neurons showed singing-modulated activity even in the absence of head movements (32/36, Table 2). These results indicate that activity of many AId neurons during song production may reflect singing-specific movements such as beak movements, respiratory actions, or gular fluttering, or other non-physical aspects of song production such as auditory-vocal feedback. Interestingly, 12 neurons showed differential response strength during head movements within singing versus non-singing periods (Figure 5B; C, right). One interpretation is that these neurons integrate information about both head movements and singing behavior, such that firing rate is enhanced specifically during the head and body movements that are performed concurrently with song; developing associations between head or postural movements and vocal behavior may be a crucial component of learning to produce female-directed song and perform courtship dance movements while singing (Morris, 1954; Balaban, 1997; Williams, 2001; Tomaszycki and Adkins-Regan, 2005).

### Additional sources of AId neuron modulation: behavioral states

As indicated above, our goal was to assess the activity of AId neurons throughout an entire session of active behaviors. As part of this approach, we devised a novel way of characterizing each recording session by classifying contiguous time periods across each session into one of five different “state” periods based on the bird’s behavior: eating, singing, active-movement, quiet-attentive, or quiescence (Fig. 6A; see Materials and Methods). Eating states were defined as periods when the bird was engaged in eating behavior, including pecking at, hulling, and ingesting seeds; although eating state periods were dominated by eating-related behaviors, other movements such as head movements or hops could also occur. Similarly, singing states included periods when birds were engaged in song production, as well as brief pauses in-between song bouts during which birds occasionally made hops, pecks, or head movements. During active-movement states, birds could produce any of the seven movements we scored as well as head and/or postural body movements that were not scored. The remaining two states characterized non-movement periods: during quiet-attentive states, the bird was alert and could make small head movements but was otherwise not moving; birds made no movements during quiescent states (quiescent states included periods from which baseline intervals were sampled in the scored-movement analyses above; see Materials and Methods).

**Figure 6.**
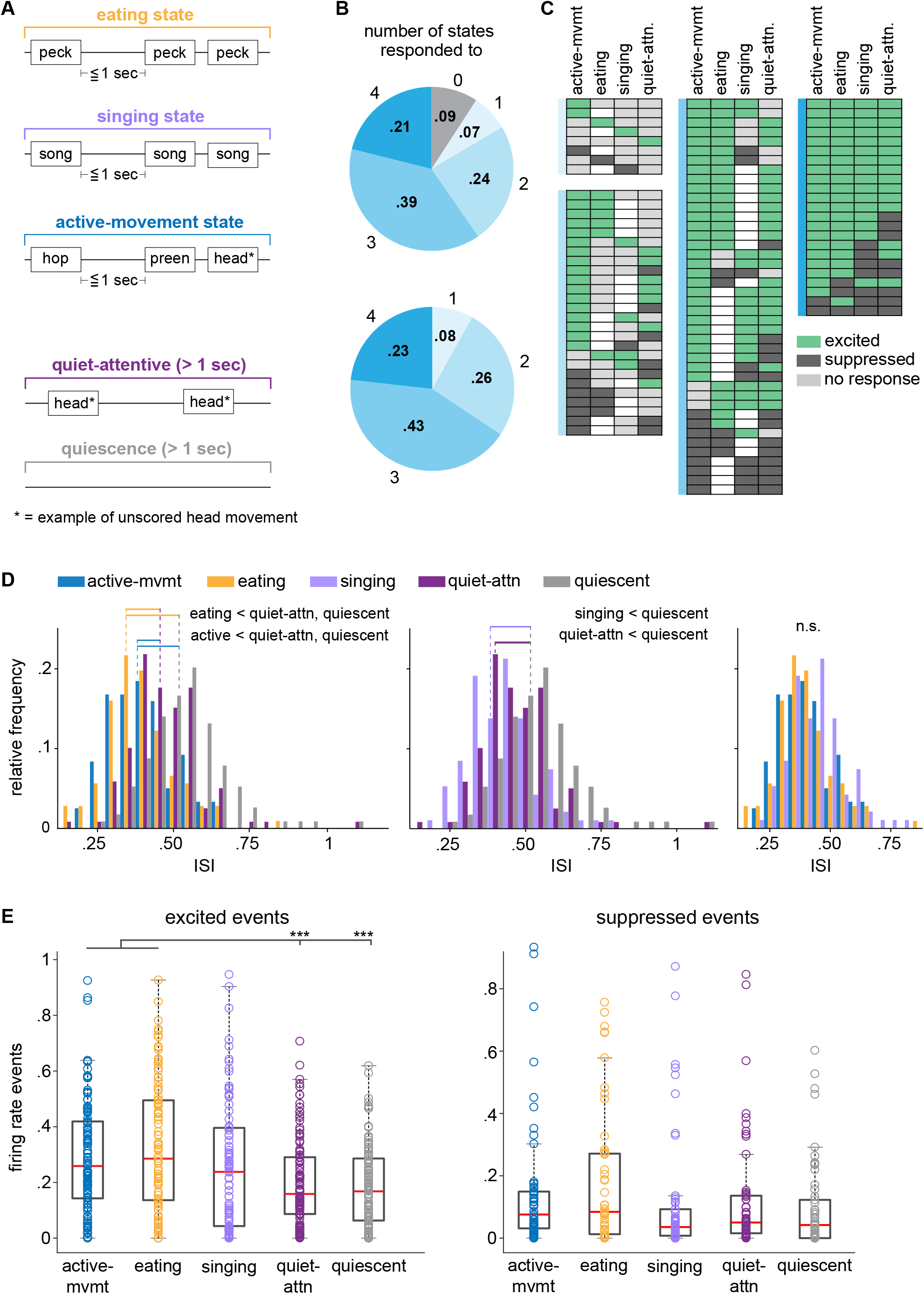
AId neurons are differentially modulated during different behavioral states. ***A***, Schematics of example eating, singing, active-movement, quiet-attentive, and quiescent states. Text boxes represent example scored and unscored (starred) movements that typically occurred within that state type, though other behaviors could also occur (see Materials and Methods). ***B***, Proportions of single AId neurons that were modulated during different numbers of state types. Top: proportions out of 109 neurons with data from quiescent state periods; 10/109 neurons were not modulated during any states, while 8/109 neurons were modulated during one state type, 26/109 during two, 42/109 during three, and 23/109 during four states. Bottom: proportions out of 99 AId neurons that were modulated during at least one state type. ***C***, Each row of each chart indicates the states during which each neuron was excited (green), suppressed (dark gray), or not responsive (light gray). Unfilled boxes indicate that no data during that state was recorded for that neuron. Charts are grouped according to colors in ***B***, based on the number of states during which neurons were modulated. ***D***, Histograms comparing distributions of inter-spike intervals (ISIs) during active-movement, eating, quiet-attentive, and quiescent states (left); singing, quiet-attentive, and quiescent states (middle); active-movement, eating, and singing states (right). Horizontal lines indicate distributions that had significantly different means; dotted lines indicate means of the respective distributions. *p*<.001 for all significant differences. ***E***, Number of excited (left) and suppressed (right) events that occurred during each state type, normalized by the total duration of each state type in a given recording session. Box-and-whisker plots indicate medians and first and third quartiles; whiskers indicate data points not considered outliers; circles represent data points from individual neurons. \*\*\**p*<.001.

For most neurons, firing rates during eating, singing, active-movement, and/or quiet-attentive state periods differed from quiescence (91%, 99/109; Fig. 6B, top; Kolmogorov-Smirnov test, p < .05). Relatively few neurons were modulated during only a single state type; rather, most neurons exhibited firing rate changes during two or more different state types, with nearly half of state-responsive neurons modulated during three different state types (Fig. 6B, bottom). Figure 6C illustrates the firing rate changes exhibited by neurons during each non-quiescent state type, categorized by the number of states during which neurons were modulated. AId neurons could show increased or decreased firing rates during non-quiescent states, and 29 out of 109 neurons were excited during one state type and suppressed during another. However, whereas single neurons were equally likely to be suppressed as excited during different discrete movements (Fig. 2B), modulation across entire state periods tended to be excitatory: within each state type, the proportion of neurons that were excited during that state was significantly greater than those that were suppressed (binomial test, active-movement and eating states p < .0001, singing and quiet-attentive states p < .05), and the overall proportion of excited state responses was greater than suppressed state responses (binomial test, p < .0001).

In accord with this pattern of results, the average inter-spike interval (ISI) during all non-quiescent states was each shorter than quiescence (Fig. 6D, left, middle; pairwise linear contrasts, Benjamini-Hochberg corrected, p < .001 in all cases). In addition, ISI’s during active-movement and eating states were shorter than during quiet-attentive states (Fig. 6D, left; p < .001 in both cases) and did not differ from singing states (Fig. 6D, right; p > .05 in both cases). Thus, although a large proportion of neurons were significantly excited during quiet-attentive states (Fig. 6C), the ISI results indicate greater increases in firing rates during periods of active behavior compared to non-movement states.

As illustrated above, activity of AId neurons in awake, behaving juveniles is both dynamic and heterogeneous during discrete movements (Figs. 1B, 2, 3). To capture these modulations during state periods, we identified excited and suppressed spiking “events”, defined as five or more contiguous 10-ms bins in which the firing rate exceeded the average firing rate during quiescent periods by ≥ 3 standard deviations (for excited events) or fell below the average quiescence firing rate by ≤ 1.5 standard deviations (for suppressed events). The average number of excited events that occurred within active-movement and eating states was each greater than during quiet-attentive and quiescence states (Fig. 6E, left; pairwise linear contrasts, Benjamini-Hochberg corrected, p < .001 in all cases). The frequency of excited events during singing states was elevated relative to quiet-attentive and quiescent states, but did not significantly differ from any of the other four states (Fig. 6E, left; pairwise linear contrasts, Benjamini-Hochberg corrected, p > .05 in all cases). The relatively modest incidence of excited events during singing compared to active-movement and eating states may indicate that firing rate modulation during singing states involves more uniform increases in tonic spike rate. The number of suppressed events did not differ between state types (Fig. 6E, right; pairwise linear contrasts, Benjamini-Hochberg corrected, p > .05 in all cases). This overall pattern of results indicates an increase in firing rate during non-quiescent states, with a greater degree of excitatory modulation during active-movement, eating and singing states expressed as shorter ISI’s relative to quiescence as well as an increase in discrete periods of higher firing during active-movement and eating states.

### Can movement-related activity fully account for modulation during behavioral state periods?

We next investigated whether firing rate increases during behavioral states could be explained by activity during the scored movements that occurred within each state. Scored behaviors occurred at different relative frequencies during different state periods: while active-movement state periods included different scored movements occurring at varying frequencies, eating and singing state periods were dominated by pecks and singing bouts, respectively (Fig. 7A). As a test of whether responses during scored movements could fully account for activity across state periods, we asked whether neurons that were excited during eating states were selectively excited during pecks, and whether neurons that were excited during singing states were selectively excited during song bouts.

**Figure 7.**
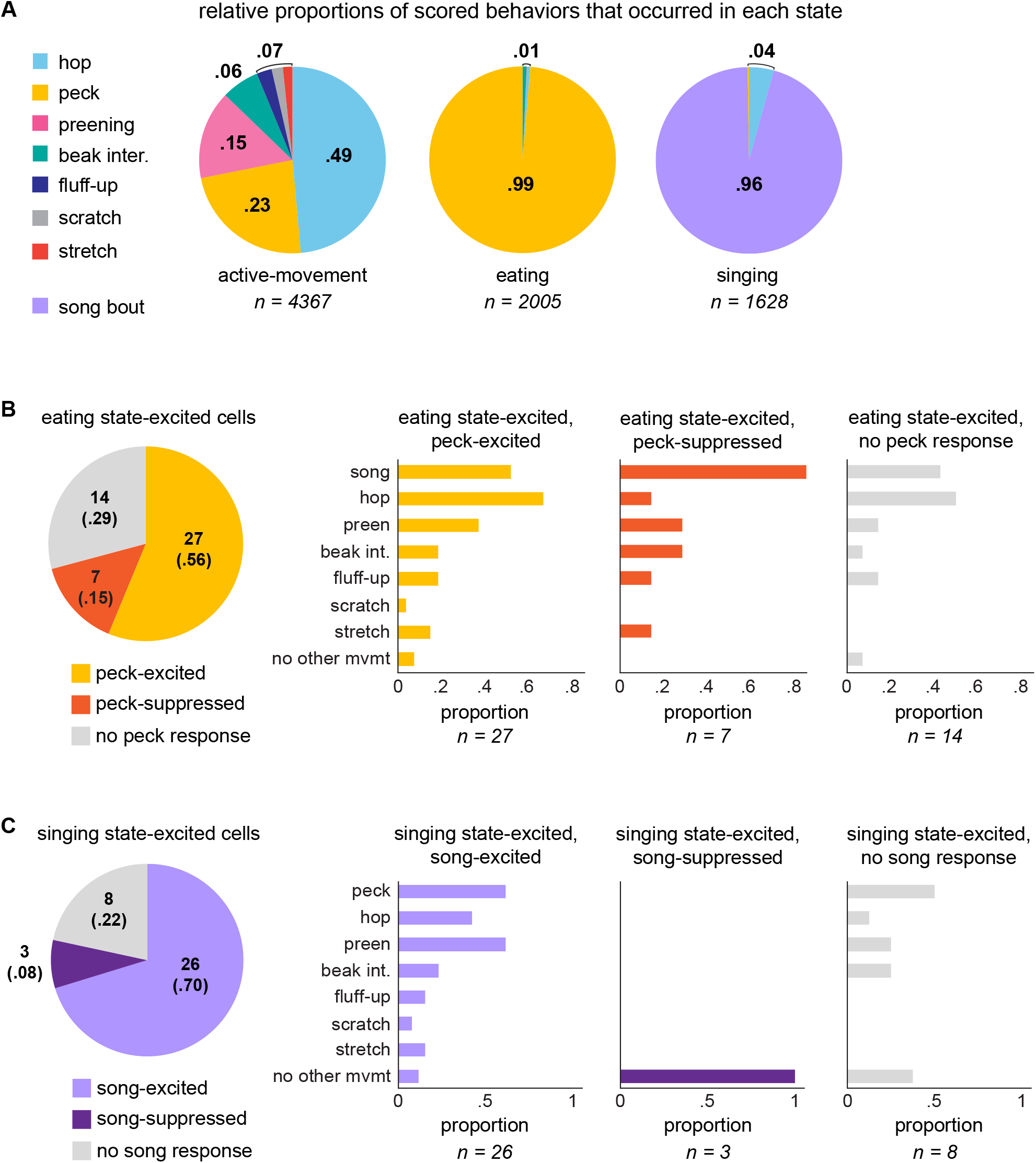
Movement-excited AId neurons are modulated by a variety of scored and unscored factors during different state periods. ***A***, Relative proportions of scored behaviors that occurred during each state type. Proportion totals include all occurrences of each of the seven scored movements as well as all song bouts. ***B***, Left: peck-responsivity of 48 neurons that were significantly excited during eating state periods. Right: proportions of eating-excited neurons that were responsive during scored behaviors, categorized by peck-responsivity: eating-excited neurons that were peck-excited (left), peck-suppressed (middle), and unresponsive during pecks (right). “No other movement” represents neurons that did not respond during any other scored behavior and were thus selectively peck-excited (left), peck-suppressed (middle), or nonresponsive during any scored movement (right). Italicized numbers indicate number of neurons within each peck-responsivity grouping. ***C***, Left: song-responsivity of 37 neurons that were significantly excited during singing state periods. Right: proportions of singing state-excited neurons that were responsive during scored movements, categorized by song-responsivity: singing state-excited neurons that were song-excited (left), song-suppressed (middle), and unresponsive during song bouts (right). “No other movement” represents neurons that did not respond during any other scored behavior and were thus selectively song-excited (left), song-suppressed (middle), or nonresponsive to any scored behavior (right). Italicized numbers indicate number of neurons within each song-responsivity grouping.

Figure 7B illustrates the movement responsivity of 48 neurons that were excited during eating states (48/99 state-responsive neurons, 48%), grouped by peck-responsivity. A majority of neurons that were excited during eating behavior were also excited during peck movements (27/48, 56%; Fig. 7B). Notably, however, these peck-excited neurons were also responsive during other scored movements – including ones that did not occur within a given state: for instance, of the 27 eating-excited neurons that were excited during pecks, 14 (52%) were also song-responsive and 10 (37%) were preening-responsive, even though these latter two behaviors never occurred during eating states (Fig. 7A, B). Likewise, most neurons that were excited during singing state periods were excited during song bouts, but not selectively so; singing state-excited neurons were also responsive during other scored movements that did not occur during singing states, such as pecking and preening (16/27, 59% each; Fig. 7C). This pattern of results supports the idea that AId neurons can be modulated by multiple factors, showing increased firing rates during a given state as well as during discrete movements that occur outside of that state type.

Although activity during scored pecking movements could explain eating state activity for most neurons, many neurons that showed excitation during eating states were not excited during pecks: 14 out of 48 (29%) neurons were excited during eating states yet were not responsive during pecks, and seven out of 48 (15%) eating-excited neurons were suppressed during pecks, indicating that the heightened firing rate of these neurons during eating did not relate to pecking behavior (Fig. 7B). Among these 21 neurons, 17 (81%) responded during at least one other movement type besides pecks. Thus, although pecks were virtually the only scored movements that occurred within eating states (Fig. 7A), neurons could show significant excitation during eating states that was not accounted for by activity during pecking movements, as well as firing rate modulation during other scored movement types that occurred outside of eating states. Increased firing rates during eating may be related to other unscored factors such as head movements or hulling behavior, or external sensory inputs. Similarly, while excitation during singing state periods could be explained by song production activity for the majority of neurons (26/37, 70%), 11 neurons were excited during singing state periods but unresponsive or suppressed during song bouts (30%; Fig. 7C). These results extend our data from movement analyses and highlight the complexity of the information that AId neurons are likely integrating: single neurons can demonstrate robust, differentially patterned responses during specific movements while also exhibiting firing rate changes during state periods that are unrelated to those movements.

While the majority of neurons we recorded were responsive during one or more scored movements, 18 out of 119 cells (15%) were not significantly modulated during any scored movement. However, most of these “movement-unresponsive” neurons were nonetheless modulated during at least one state type compared to quiescence (15/18, 83%; Fig. 8A). We grouped these unresponsive neurons by the state during which their firing rate was highest and tested whether activity was differentially modulated between state groups. The firing rate of neurons within each preferred state group tended to be higher relative to all other state types despite consisting entirely of neurons that were not responsive during any scored movement, although group differences did not reach significance due to small group sizes (Fig. 8B; pairwise linear contrasts, Benjamini-Hochberg corrected, p > .05 in all cases). This pattern of results suggests that some of the neurons unresponsive during discrete movements are nonetheless modulated by other unscored factors as the juvenile is actively behaving.

**Figure 8.**
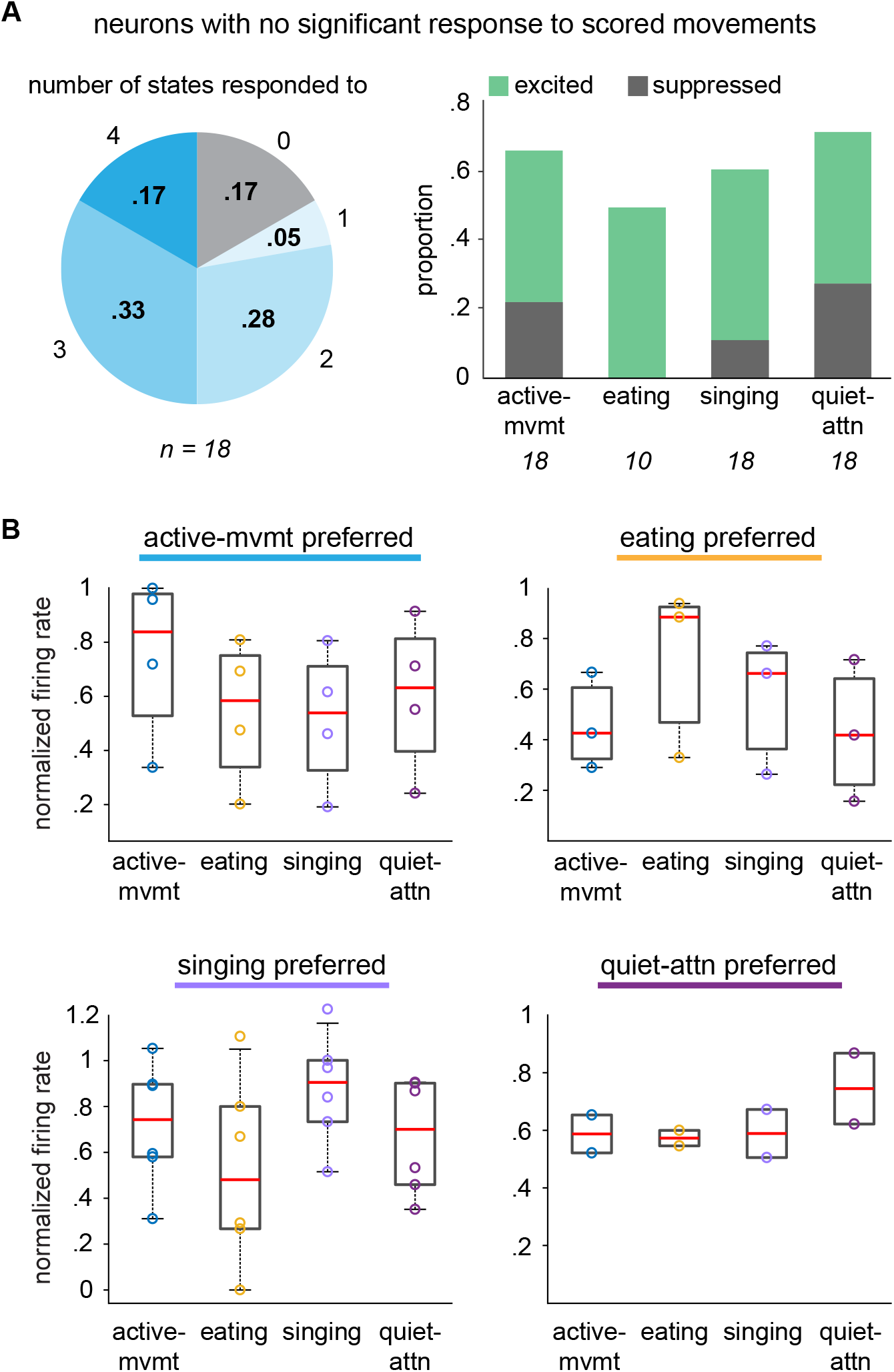
AId neurons that lack responsivity to scored movements are nonetheless modulated during different states. ***A***, Left: proportions of non-responsive AId neurons that were modulated during different numbers of state types. 3/18 neurons were not modulated during any states; 1/18 was modulated during one state type, 5/18 during two, 6/18 during three, and 3/18 during four state types. Right: proportions of non-responsive neurons that were significantly excited (green) or suppressed (dark gray) during each state type. Italicized numbers indicate number of non-responsive neurons recorded during each state type. ***B***, Mean firing rates of neurons that were not responsive during any scored movement during each state type, classified by neurons that showed maximal average firing rate during active-movement (top left), eating (top right), singing (bottom left), and quiet-attentive (bottom right) states. Firing rates within each state type were normalized by the standard deviation of each neuron’s firing rate across the recording session. Box-and-whisker plots indicate medians and first and third quartiles; whiskers indicate data points not considered outliers; circles represent data points from individual neurons.

## DISCUSSION

### Heterogeneous activity within AId suggests multi-dimensional tuning including both sensory and motor components

We found that most AId neurons were selective for single movements or for different combinations of movements. Neurons responsive during different movements frequently demonstrated excitation during one movement and suppression during another. Moreover, individual responses could include both transient excitation at movement onset and/or offset as well as suppression of average firing rate during the movement itself, or vice versa. The diversity of neural responses in AId is strikingly similar to the response profile of neurons in macaque motor cortex, where single neurons demonstrate heterogeneous, multiphasic temporal patterns of activity across reaching movements (Churchland and Shenoy, 2007). Such complex responses may result from multiple inputs relating to different movements or aspects of movements onto single AId neurons, as well as local transformation of afferent inputs. AId includes a local inhibitory network, evidenced by the fact that blocking GABA-A receptors in AId of anesthetized zebra finches elicits increased spontaneous firing rates, and parvalbumin expression is higher in AId compared to surrounding motor cortex (Mello et al., 2019; Yuan and Bottjer, 2019). Similarly, mammalian motor cortex contains a substantial population of inhibitory interneurons, which have been implicated in both regulating plasticity during motor skill learning and coordinating activity across motor cortex during behavior (Jacobs and Donoghue, 1991; Hess and Donoghue, 1994; Hess et al., 1996; Markram et al., 2004; Stagg et al., 2011; Donato et al., 2013; Chen et al., 2015; Kida et al., 2016; Adler et al., 2019). Studies have suggested that blocking inhibition unmasks latent excitatory connections between spatially distant motor cortical neurons, providing a mechanism by which dynamic modulation of inhibition could flexibly reorganize connectivity and coordinate population activity across motor cortex (Jacobs and Donoghue, 1991; Spiro et al., 1999; Schneider et al., 2002; Capaday, 2004). AId receives topographic input from parallel circuits that process auditory, visual, and somatosensory information, so a similar mechanism to link different neuronal subpopulations within AId would be advantageous for facilitating sensorimotor integration across modalities (Zeier and Karten, 1971; Bottjer et al., 2000; Paterson and Bottjer, 2017).

The heterogeneous response profile of motor cortical neurons across taxa raises interesting questions about what factors contribute to the tuning of these neurons. Modulation of motor cortical activity has been associated with a variety of behavioral parameters in arm-reaching tasks, including movement direction, speed, trajectory, limb position, and joint angle (Evarts, 1968; Cheney and Fetz, 1980; Georgopoulos et al., 1982, 1988; Schwartz et al., 1988; Fu et al., 1993; Schwartz and Moran, 2000; Reina et al., 2001; Paninski et al., 2004; Churchland and Shenoy, 2007; Hatsopoulos et al., 2007). Increasing evidence indicates that multiple parameters can be reflected in the activity of single neurons, suggesting that highly integrated multi-modal tuning may be a fundamental feature of motor cortical activity. For instance, recordings from macaques during unrestrained arm movements showed that parameters such as movement direction or end position of the limb could account for only a portion of spiking patterns in single motor cortical neurons, indicating that individual neurons may be tuned in a multidimensional space and that testing neural activity relative to any single parameter may account only partially for multidimensional tuning profiles (Ashe and Georgopoulos, 1994; Fu et al., 1995; Moran and Schwartz, 1999; Aflalo and Graziano, 2006, 2007). Likewise, we found that single AId neurons could be modulated both during individual movements and during behavioral state periods that did not include those movements, indicating that single neurons were modulated by multiple factors.

Given convergent input from a diverse array of multi-modal processing streams, the tuning profiles of neurons in both avian and mammalian motor cortex often include not only motor but also sensory responsivity. For instance, neurons in RA, which lies adjacent to AId in songbird motor cortex, drive vocal motor output in singing birds and demonstrate robust responses to playback of song stimuli in anesthetized birds (Nottebohm et al., 1976; Doupe and Konishi, 1991; Vicario and Yohay, 1993; Wild, 1993; Yu and Margoliash, 1996; Dave et al., 1998; Dave and Margoliash, 2000; Leonardo and Fee, 2005; Kojima and Doupe, 2007; Sober et al., 2008; Yuan and Bottjer, 2019). Recordings from macaque motor cortex have likewise demonstrated sensitivity to non-motor stimuli: for instance, in visually-guided target-reaching paradigms, some motor cortical neurons exhibit selective activity related to the visual target, regardless of the limb trajectory used to reach that target (Tanji and Evarts, 1976; Evarts and Fromm, 1977; Murata et al., 1997; Shen and Alexander, 1997). The activity of AId neurons may be similarly modulated by integration of a variety of motor and sensory factors to produce heterogeneous responses with complex temporal patterning during diverse movements. In further support of this idea, we also found neurons whose activity was modulated across behavioral state periods but not during any scored movements, suggesting that activity in these neurons may relate to non-motor factors such as visual or attentional processing (Knudsen et al., 1995; Winkowski and Knudsen, 2007; Fernández et al., 2020).

### Multi-modal integration provides behavioral context for voluntary movements

We found that most AId neurons demonstrated altered response strength during movements relative to quiescence; under the conditions of our recordings, firing rate modulations occurred most often during pecks, preening, and/or hops. It is difficult to know whether movement-related activity in AId neurons is pre-motor, modulatory, or reflective of movement feedback or external sensory inputs. AId neurons project to several targets, including the striatum, a dorsal thalamic zone, the lateral hypothalamus, a thalamic nucleus that relays to cerebellum (SpM, medial spiriform nucleus), deep layers of the tectum, broad areas of the pontine and midbrain reticular formation, and the ventral tegmental area (Fig. 1A) (Bottjer et al., 2000). The medial pontine reticular formation contains premotor neurons that contribute to neck and locomotive movements in other avian species (Steeves et al., 1987; Valenzuela et al., 1990; Dubbeldam, 1998; Wild and Krützfeldt, 2012); peck, preening, and hop-related activity may in part reflect these projections to premotor centers. Moreover, previous studies have found evidence of induced ZENK activity in AId specifically during hopping behavior (Feenders et al., 2008).

However, both prior evidence and data presented here suggest that movement-related activity in AId is unlikely to directly drive peck, preening, and/or hopping behavior per se. Importantly, lesions of AId in juvenile birds do not disrupt song output or induce noticeable motor deficits, suggesting that AId neurons are not driving voluntary pecking or hopping movements (Bottjer and Altenau, 2010) [cf. Mandelblat-Cerf et al., 2014, in which AId-targeted lesions may have extended laterally beyond LMAN-shell-recipient AId to encompass more lateral regions of intermediate arcopallium, resulting in motor impairment (Wild and Farabaugh, 1996; Wild and Krützfeldt, 2012).] Moreover, the results presented here demonstrate that single AId neurons do not respond consistently during one particular type of movement. For instance, we found a substantial population of peck-responsive neurons that modulated their firing rate during pecks when the bird was eating but not when the bird pecked at other objects around the cage (or vice versa), even though pecking movements in these different contexts would recruit many of the same muscle groups. Furthermore, behavioral and functional experiments across avian species have implicated intermediate arcopallium in highly integrative, complex behaviors that extend beyond pure motor control, including ingestive behaviors, working memory processing, fear conditioning, and vocal learning (Lowndes and Davies, 1994; Lowndes et al., 1994; Knudsen and Knudsen, 1996; Aoki et al., 2006; Campanella et al., 2009; Saint-Dizier et al., 2009; Bottjer and Altenau, 2010; Achiro et al., 2017). One such example is a region comprising caudal arco- and nidopallium, which partially overlaps with AId and exhibited increased 2-deoxyglucose uptake when adult male zebra finches participated in their first courtship experience following several weeks of isolation from females; the amount of glucose consumption in this region correlated positively with isolation time but not with amount of movement activity (Bischof and Herrmann, 1986).

Rather than generically driving motor behavior per se, an important function of motor cortical circuitry is incorporating sensory information to appropriately direct motor output based on an animal’s environment and/or goals; this sensorimotor integration is necessary for voluntary movements such as goal-directed motor behaviors (e.g., object-directed grasping) as well as adaptive movements based on environmental perturbations. Avian and mammalian motor cortices receive multi-modal inputs and target brainstem regions, making them ideally situated to carry out sensorimotor integration during voluntary behaviors. In macaques, neurons in motor cortical areas demonstrate a selective response when grasping at a particular object and corresponding visual selectivity for the same object when the monkey fixates on the object without grasping; inactivation of the same motor cortical region resulted in grasping deficits due to disrupted preparatory hand shaping that was inappropriate for the target object, suggesting a specific impairment in visuomotor transformations for targeted grasping rather than a gross motor impairment of hand movements (Murata et al., 1997; Fogassi, 2001; Rizzolatti and Luppino, 2001). Some AId neurons may serve a similar function in integrating sensory information to provide appropriate context for behavior – for example, the “eating-peck” responses observed here could represent an integrated response when the visual stimulus of seed is present as the bird pecks; these neurons could link visual information about seed with somatosensory information to contribute specifically to food-directed pecking behavior. In contrast, neurons that showed excitation during pecks associated with non-eating behaviors may process diverse inputs in the context of object exploration. Notably, AId neurons could provide appropriate context for voluntary behaviors without directly driving motor actions – for instance, rather than directly linking sensory information to premotor centers, some neurons could instead feed multi-modal information back into ascending reticular or tectal pathways to contribute to goal-directed behaviors, or differentially ascribe value to environmental cues depending on the animal’s current state or needs (Burgess et al., 2018).

### AId is uniquely situated to mediate learning and performance of both vocal and non-vocal elements of song behavior

Although motor cortex is generally well-situated to integrate multi-modal information related to a variety of goal-directed movements, AId’s unique connections suggest it may occupy a specific role in vocal learning and behavior. LMAN-CORE neurons that drive vocal motor output in juvenile birds make robust collateral projections into AId at 20-35 dph that substantially decline by 45 dph (Miller-Sims and Bottjer, 2012; unpublished observations). While our data set did not include ages young enough to test the functional role of this connection between LMAN-CORE and AId, information from this developmentally regulated projection may play a critical role during the earliest stages of vocal practice and influence patterns of connectivity within AId that contribute to sensorimotor processing during subsequent learning. In addition, AId projects to higher-order thalamic nuclei that are linked to vocal learning, DLM (dorsolateral nucleus of the medial thalamus) and DMP (dorsomedial nucleus of the posterior thalamus) (Fig. 1A) (Bottjer et al., 2000). DLM is required for normal song behavior and projects to LMAN-shell, whereas DMP projects to MMAN (medial magnocellular nucleus of anterior nidopallium); both LMAN and MMAN are required for development of an accurate imitation of tutor song (Bottjer et al., 1984; Foster et al., 1997; Vates et al., 1997; Foster and Bottjer, 2001; Aronov et al., 2008; Goldberg and Fee, 2011; Chen et al., 2014). AId also projects to lateral hypothalamus, striatum, and the area of dopaminergic neurons in the ventral tegmental area that projects to a nucleus in avian basal ganglia necessary for song learning (Fig. 1A) (Bottjer et al., 2000); these limbic-related projections further suggest that AId neurons are well situated to contribute to vocal learning and behavior.

One hypothesis to integrate these unique connections with the multi-modal integrative function of motor cortex is that some AId neurons may be involved in mediating learning and performance of movements in the context of song behavior. Song production in zebra finches is a courtship behavior during which males vocalize while performing a dance-like sequence of hopping movements oriented towards a female (Morris, 1954; Williams, 2001; Cooper and Goller, 2004; Dalziell et al., 2013; Ota et al., 2015; Ullrich et al., 2016); the temporal patterning of dance movements during song production is significantly correlated between father-son pairs of zebra finches, suggesting that non-vocal behaviors that accompany singing may be learned as well (Williams, 2001). Establishing a social context for courtship behavior likely involves sensory cues. For instance, adult males can use visual cues to select between two female birds shown in a silent video feed, but the addition of auditory cues induces stronger courtship responses (Galoch and Bischof, 2006, 2007). AId neurons are ideally positioned to integrate environmental context cues when females are present to guide learning and performance of movements that accompany singing.

In this framework, the relatively high proportions of peck, preening, and hopping-related responses observed here may reflect the fact that beak movements and hopping are important components of courtship behavior. In quail-chick chimeras, chicks that received transplants of lower brainstem somites from quails retained chick-like call structures but adopted quail-like patterns of head movements specifically during vocalizations; similar head movements made outside of vocalization periods were not affected, indicating the presence of specialized circuitry that mediates movements in the context of vocal behavior (Balaban, 1997). Circuitry processing non-vocal elements of singing in zebra finches may be similarly specialized; our results suggest that investigating how activity patterns during singing correspond to movements of different peripheral targets would benefit hypotheses for mechanisms of song production: for instance, we found a subpopulation of AId neurons that exhibited differential response strength during head movements that occurred during singing versus non-singing periods. These responses may indicate integration between non-vocal and vocal elements of song behavior, such that neural activity is enhanced specifically during head movements that accompany singing. This hypothesis raises an interesting prediction: although AId lesions in juvenile birds do not induce any gross motor deficits, it is possible that hopping and/or head movements performed during song production would be disrupted. Such a result would be consistent with previous studies in which c-Fos expression in AId was increased after adult male zebra finches performed non-singing courtship behaviors directed towards a live female (Kimpo and Doupe, 1997). Involvement of AId in courtship-related movements would draw an interesting parallel to mammalian studies that have suggested that motor cortex is parceled into “action zones” that each process information for different ethologically relevant categories of movement (Graziano, 2006; Graziano and Aflalo, 2007). For instance, stimulation of one region of macaque motor cortex results in the animal closing its hand in a grip while bringing it to its mouth and opening its mouth, as if eating an object, while stimulation of another region results in the monkey raising its arm and turning its head sharply to one side as if in defense (Graziano et al., 2002). In this context, adjacent motor cortical regions RA and AId could serve as an “action zone” that mediates the vocal and non-vocal movements that comprise song behavior.

While brainstem projections may mediate sensorimotor processing during vocal motor performance, the thalamic and midbrain projections of AId that give rise to recurrent feedback loops through cortico-basal ganglia circuitry may integrate multi-modal information to facilitate song learning. Although AId does not drive song output (Bottjer et al., 2000; Bottjer and Altenau, 2010), we found a substantial population of singing-responsive neurons. For a large proportion of these neurons, the changes in firing rate during singing could not be attributed to any of the movements that we scored (including head movements). These firing rate modulations may instead be related to singing-specific movements such as beak movements or gular fluttering, or sensory activity such as auditory or proprioceptive feedback (Goller et al., 2004; Ohms et al., 2010; Bottjer and To, 2012; Riede et al., 2013). Alternatively, singing-related activity could reflect active evaluations of the juvenile’s vocal behavior during sensorimotor practice – iterative evaluations between self-generated output and the goal tutor song are essential for guiding accurate refinement of the juvenile’s song, and evidence of neural activity processing these comparisons has been reported in LMAN-shell, which projects directly to AId (Achiro et al., 2017). Importantly, successful song learning also requires multiple factors beyond simply matching vocal output to an auditory goal – for instance, vocal learning in juvenile zebra finches that are tutored with only passive playback of the tutor song is severely impaired, whereas pairing auditory tutoring with a visual model of an adult zebra finch enhances learning (Derégnaucourt et al., 2013; Ljubičić et al., 2016). Moreover, visual cues provided during singing, such as wing strokes or fluff-ups from adult females, provide feedback that can influence juvenile vocal learning (West and King, 1988; King et al., 2005; Carouso-Peck and Goldstein, 2019). Multi-modal inputs from dNCL and singing-related inputs from LMAN may converge in AId, integrating important non-vocal and vocal elements of courtship song behavior that must be learned during a sensitive period of development.

---

**Video 1**. Example video of neural activity recorded on a single channel while the juvenile zebra finch hopped around the cage, demonstrating firing rate increases whenever the bird hopped towards the left side of the cage. Vertical lines above the raw activity trace indicate spikes from a single neuron sorted from the extracellular activity.

